# A Capped Tudor Domain within a Core Subunit of the Sin3L/Rpd3L Histone Deacetylase Complex Binds Nucleic Acids

**DOI:** 10.1101/2021.08.09.455673

**Authors:** Ryan Dale Marcum, Joseph Hsieh, Maksim Giljen, Yongbo Zhang, Ishwar Radhakrishnan

**Author notes:** Contact information for coauthors.

## Abstract

Chromatin-modifying complexes containing histone deacetylase (HDAC) activities play critical roles in the regulation of gene transcription in eukaryotes. These complexes are thought to lack intrinsic DNA-binding activity, but according to a well-established paradigm, they are recruited via protein-protein interactions by gene-specific transcription factors and post-translational histone modifications to their sites of action on the genome. The mammalian Sin3L/Rpd3L complex, comprising more than a dozen different polypeptides, is an ancient HDAC complex found in diverse eukaryotes. The subunits of this complex harbor conserved domains and motifs of unknown structure and function. Here we show that Sds3, a constitutively associated subunit critical for the proper functioning of the complex, harbors a type of Tudor domain that we designate the capped Tudor domain (CTD). Unlike canonical Tudor domains that bind modified histones, the Sds3 CTD binds to nucleic acids that can form higher-order structures such as G-quadruplexes, and shares similarities with the knotted Tudor domain of the Esa1 histone acetyltransferase (HAT) that was previously shown to bind single-stranded RNA. Our findings expand the range of macromolecules capable of recruiting the Sin3L/Rpd3L complex and draws attention to potentially new roles for this HDAC complex in transcription biology.

## Introduction

Post-translational modifications of core histones constitute a common molecular mechanism for regulating transcription by modulating DNA template accessibility to RNA polymerases, regulatory factors, and other effectors (1,2). Among various post-translational modifications, acetylation of lysine residues is not only abundant but also one that is characterized by high turnover, consistent with its central role in the dynamic induction and repression of genes (3). Deacetylation of histones in mammals is mediated in large part by histone deacetylases (HDACs) 1, 2, and 3 (4-6). These enzymes, found in at least six giant multiprotein complexes including the Sin3L/Rpd3L, Sin3S/Rpd3S, NurD, LSD1-CoREST, MiDAC, and SMART/NCoR complexes (7-11), exert their effects following recruitment to specific sites on the genome by DNA-bound transcription factors and/or specific histone modifications.

The Sin3L/Rpd3L complex is the prototypical HDAC complex found in organisms as diverse as yeast and human (8,12). The complex plays fundamental roles in mammalian biology, regulating a wide array of genes involved in the cell cycle, differentiation, metabolism, and stem cell maintenance (13-15). The 1.2-2 MDa mammalian complex harbors at least 10 constitutively associated subunits including Sin3A/B, HDAC1/2, RBBP4/7, Sds3/BRMS1/BRMS1L, SAP30/SAP30L, ING1b/ING2, SAP130a/b, ARID4A/B, FAM60A, and SAP25 (paralogous proteins in this list are separated by a ‘/’). The first five subunits on the list comprise the core complex because of their essential roles in complex assembly and stability (16-19); these subunits along with the ING subunits have orthologs in yeast. Whereas the RBBP, ING, and ARID4 subunits harbor WD-40, PHD, and Royal Family domains that bind unmodified and modified histones, the other subunits of the complex harbor conserved domains of unknown structure and function.

In the course of our studies to define the molecular roles of the key subunits of the Rpd3L/Sin3L complex, we previously described a novel zinc finger motif shared by the SAP30 and SAP30L subunits of this complex that we later showed turbocharges HDAC activity in response to small-molecule effectors such as inositol phosphates derived from membrane lipids (20,21). We also showed how the Sds3 subunit provides a dimerization function for the complex that involves a region that assembles into a two-stranded antiparallel coiled-coil helix (22). We further showed that the subunit plays a critical role in core complex assembly by engaging directly and independently with Sin3 and HDACs; the subunit and its paralogs have been implicated in interactions with other subunits of the complex as well as with sequence-specific DNA transcription factors (22-24). At the cellular level, the subunit plays a critical role in the proper segregation of chromosomes during cell division by targeting the Sin3L/Rpd3L complex to pericentric heterochromatin (16). Recently, the subunit has also been implicated in resolving co-transcriptionally generated R-loops (featuring RNA-DNA hybrids and single-stranded DNA) that are a major source of genomic instability (25).

Two paralogs of Sds3 have been described including the breast cancer metastasis suppressor 1 (BRMS1) and a BRMS1-like protein called BRMS1L that share many domains found in Sds3 (24,26). All three proteins are found in Sin3L/Rpd3L complexes and share certain key structural and functional features but are not functionally redundant. For example, disruption and downregulation of BRMS1 and BRMS1L is associated with metastasis of multiple types of cancers, whereas overexpression suppresses this effect through a mechanism involving repression of several metastasis-associated protein-coding and microRNA genes (27-31). However, Sds3 overexpression fails to compensate for *BRMS1* deletion or epigenetic silencing in breast cancer and does not suppress metastasis (32); the molecular basis for this observation remains obscure (33,34), warranting deeper structural and functional studies to understand the molecular roles of Sds3 and its paralogs.

Here, we describe the structure of another conserved domain of an unknown function in the Sds3 subunit that is shared with one of its paralogs, BRMS1L, but not with BRMS1. Our structural and biochemical analyses suggest that the domain broadly shares an SH3-like β-barrel fold found within many chromatin-binding transcription factors but instead of binding chromatin, the domain binds nucleic acids including both RNA and DNA.

## Results

### A Conserved C-terminal Domain (CTD) of Unknown Structure & Function in Sds3 and BRMS1L

Sequence analysis of Sds3 and BRMS1L orthologs from human to zebrafish revealed an ∼80 residue region at the C-termini of the respective proteins with a pattern of conservation that suggested a well-conserved, independently folded domain (**Figure 1a**); the next well-conserved segment corresponding to the previously characterized Sin3-interaction domain (SID) resides ∼30 residues N-terminal of this domain (22). Searches conducted using the mouse Sds3 protein of the RCSB PDB database using BLAST as well as the repository of templates in the SWISS-MODEL homology modeling server failed to identify any *bona fide* structural homologs for this domain (*i*.*e*., no suitable templates in the so-called safe zone for homology modeling).

**Figure 1.**
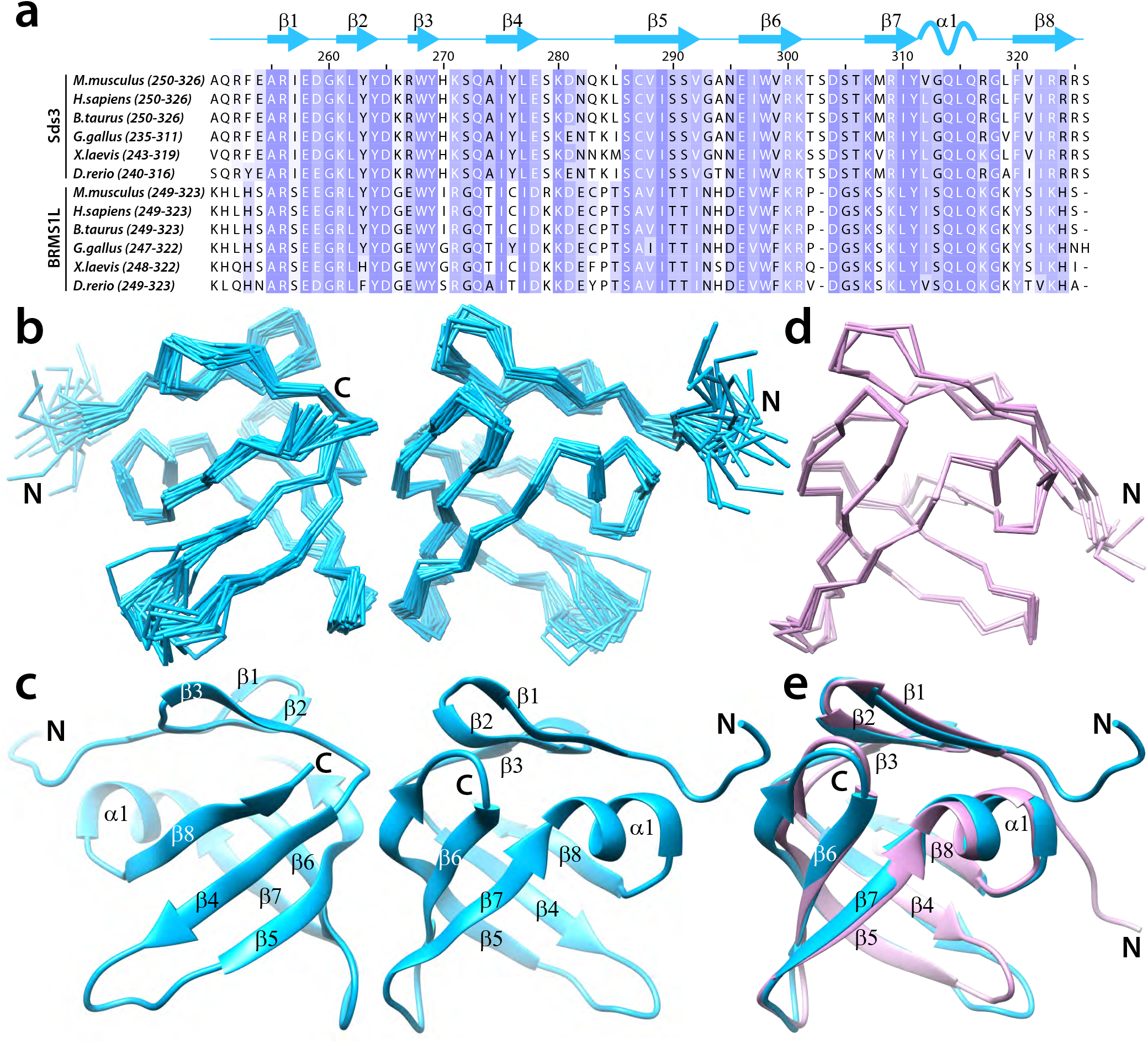
Solution structure of Sds3 CTD and comparison with prediction. (**a**) A CLUSTAL *Ω*-guided multiple sequence alignment of a ∼80-residue region at the C-terminus of Sds3 and BRMS1L orthologs from various eukaryotes. Two views differing by a 180° rotation around the vertical axis of (**b**) a best-fit backbone superposition of the ensemble of 20 conformers and (**c**) the corresponding representative structure. (**d**) A best-fit backbone superposition of the top five solutions returned by Rosetta. (**e**) A best-fit backbone superposition of the representative NMR structure with the highest-ranked Rosetta prediction. The cartoon on top of panel **a** identifies the locations of various secondary structural elements in the solution structure.

### Sds3 CTD Adopts a Unique Variation of a Common Fold

To gain insights into the structure and function of the CTD, we expressed and purified a recombinant protein corresponding to residues 250-326 of mouse Sds3. A ^1^H-^15^N correlated NMR spectrum of this protein was characterized by narrow and well-dispersed resonances indicative of a folded domain (**Supplementary Figure S1**). The solution NMR structure of the domain was determined using a combination of ^1^H-^1^H NOE-based distance and backbone chemical shift-based torsion angle restraints (**Table 1**). Structure determination and refinement resulted in an ensemble of 20 converged conformers with average RMSDs in ordered regions of 0.52 Å and good agreement with experimental restraints and excellent backbone and covalent geometry (**Figure 1b; Table 1**). The domain comprises eight strands and a short helix (**Figure 1c**). Except at the N- and C-termini and the loop connecting β4 and β5, the conformers adopt highly similar backbone conformations. Strands β4 to β8 form an antiparallel five-stranded closed β-barrel fold, reminiscent of SH3-domains, with one mouth of the barrel capped by the three-stranded β-sheet formed by β1, β2, and β3.

**Table 1.**
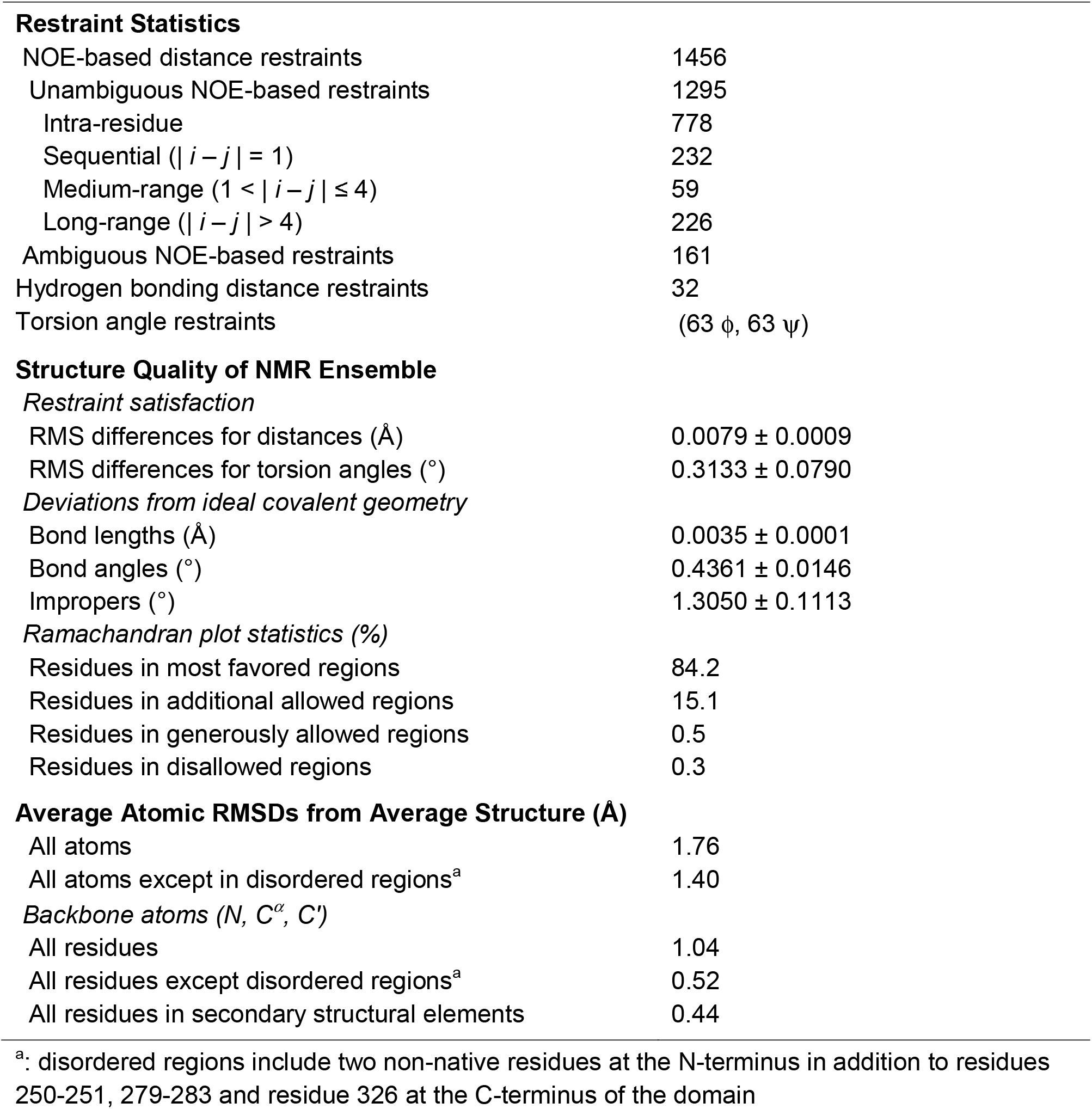
NMR Structure Determination Statistics

To test the reliability of predictions by *de novo* methods, the mouse Sds3 sequence was submitted to the Robetta server for tertiary structure prediction using TrRefineRosetta (35). The top five solutions returned by this method all had the same overall fold (**Figure 1d**), and remarkably, the backbone RMSD for the 65 C^*α*^ atom pairs involved in the best-fit superposition between the top solution and the representative structure from the NMR ensemble was 0.84 Å (**Figure 1e**). Thus, although homology modeling (aka comparative modeling) methods failed to detect a suitable template for modeling, Rosetta could readily and reliably predict the structure of this domain.

To gain insights into the domain’s function, we sought to establish the closest homolog at the structural level by searching the RCSB PDB database using DALI (36). Although DALI returned many hits, the one in the PDB25 database with the highest Z-score (6.9) corresponded to the bromo adjacent homology (BAH) domain of the *Zea mays* protein ZMET2, a DNA cytosine-5-methyltransferase ((37); **Figure 2a**). A best-fit superposition of all the regular secondary structural elements shared by the two proteins, except for the slightly elongated helical segment in Sds3, yielded a backbone RMSD of 1.87 Å. The ZMET2 BAH domain binds to histone H3K9me_2_, in a manner reminiscent of methyllysine/methylarginine-binding by the so-called Royal domains including the Tudor, MBT, chromobarrel, and PWWP domains ((38); **Supplementary Figure S2**). Comparisons with a representative structure for each of these four members of the Royal family suggested that the Sds3 CTD might be evolutionarily closest to the Tudor domain with a best-fit superposition of backbone atoms of 1.39 Å (**Figure 2b**). The Sds3 CTD harbors two features that distinguish it from regular Tudor domains: a longer helical segment linking the two C-terminal strands of the β-barrel and a three-stranded β-sheet at the N-terminus that closes one edge of the barrel, thereby capping it. Therefore, we refer to the Sds3 domain as a ‘capped’ Tudor domain (CTD).

**Figure 2.**
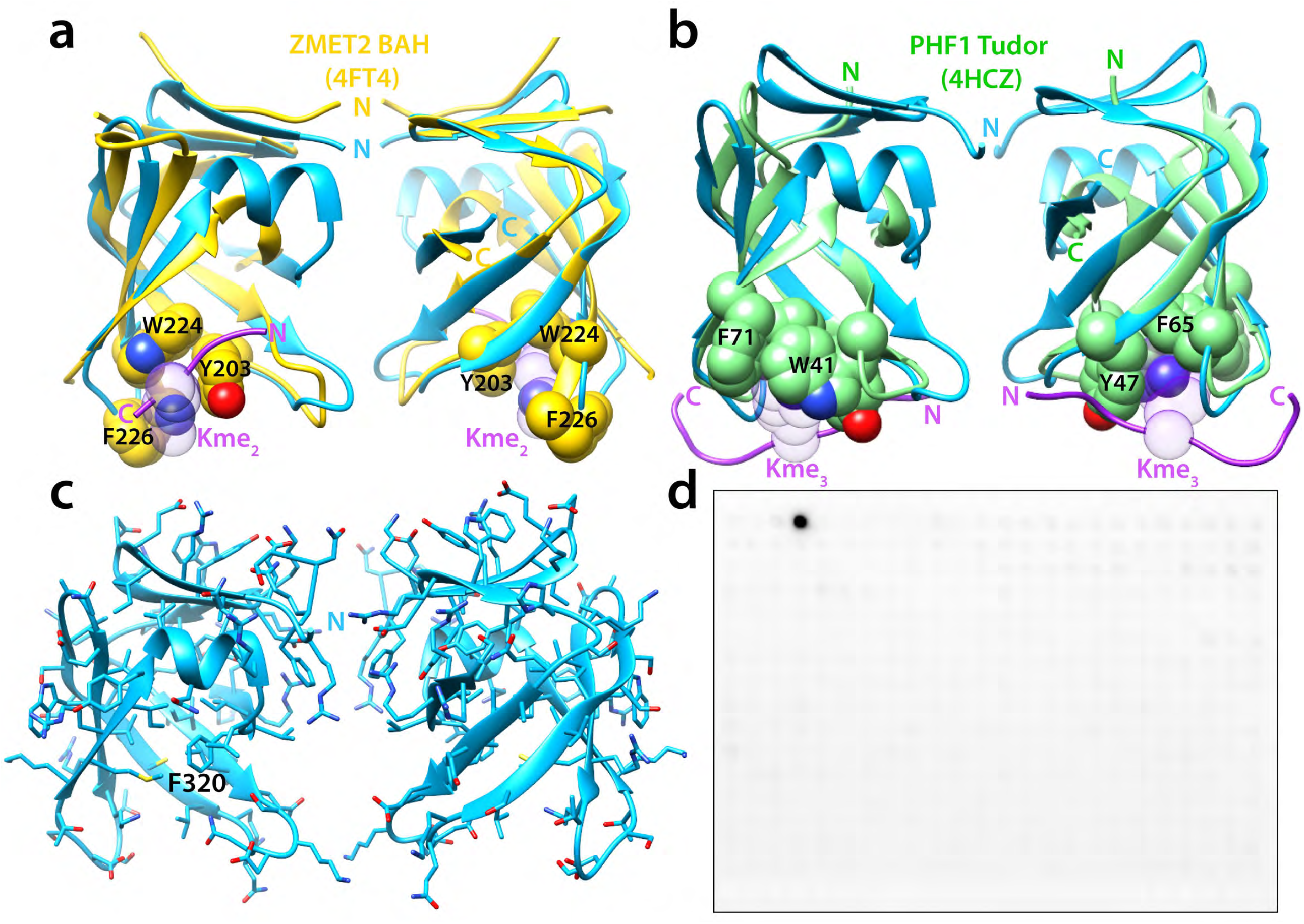
Structural relatives and insights into a potential function for the Sds3 CTD. (**a**) A best-fit backbone superposition of Sds3 CTD (blue) with its relative ZMET2 BAH domain (yellow) as determined by DALI. The BAH domain binds to a dimethyllysine-containing histone peptide with the modified residue (rendered transparently) binding to a ‘cage’ formed by the side chains of three aromatic residues (shown in spacefilling representation). The BAH domain features two long insertions in the loop regions of Sds3 CTD connecting the β1 and β2 strands and the β6 and β7 strands; neither of them is shown for clarity. (**b**) A best-fit backbone superposition of Sds3 CTD (blue) with the PHF1 Tudor domain (green) that binds to a trimethyllysine-containing histone peptide. The modified lysine is rendered transparently while the aromatic side chains of the four residues forming the ‘cage’ are rendered in spacefilling mode. (**c**) Two views, identical to those shown in panels **a** and **b**, of the Sds3 CTD with the side chains shown in stick representation to illustrate the general lack of aromatic side chains (except for F320) on one edge of the barrel that constitutes the canonical binding pocket for modified lysines and arginines. (**d**) A binding screen conducted with purified His_6_-tagged Sds3 CTD and the MODified™ histone peptide array. The results illustrate a complete lack of histone-binding activity for the CTD. The sole dark spot in the array corresponds to a Myc peptide, included in the array as a positive control, detected by an anti-Myc antibody.

### Sds3 CTD is not a Histone-binding Module

Unlike the aforementioned BAH and Royal family domains, the Sds3 capped Tudor domain lacks an aromatic cage near the remaining edge of the β-barrel for binding methyllysine residues (**Figures 2a-2c** and **Supplementary Figure S2**). Indeed, the domain in this region is devoid of aromatic residues barring one (Phe320; **Figure 2c**). To test whether the domain could bind to post-translationally modified histones, we screened the MODified™ histone peptide array using purified protein. Although an anti-Myc antibody bound to the Myc peptide that was included in the array as a positive control, no binding was detected for Sds3 to any of the histone peptides, both modified and unmodified, in the array (**Figure 2d**). To complement these findings, NMR titrations of ^15^N-labeled Sds3 CTD were conducted with dimethyllysine, trimethyllysine, acetyllysine, and dimethylarginine as well as with an unmodified histone H3 peptide (residues 1-42). However, none of these compounds produced any discernible perturbations in the NMR spectra. Collectively, these results suggest that the Sds3 CTD has no apparent histone-binding activity.

### Sds3 CTD Surface Properties Suggest a Role in Nucleic Acid Binding

To gain clues into its molecular function, we then analyzed the surface properties of the Sds3 capped Tudor domain. Given the considerably high levels of sequence conservation (**Figure 1a**), mapping the information onto the molecular surface was not especially insightful. However, since Tudor domains have been implicated in functions other than histone binding, such as nucleic acid binding, we calculated the electrostatic potential using APBS (39) and mapped it onto the molecular surface of the domain. Doing so revealed an overwhelmingly electropositive or neutral surface with multiple, discrete patches that were strongly electropositive (**Figure 3a**). Since the Sin3L/Rpd3L complex functions, in part, by directly engaging with nucleosomes (40), we first asked whether the domain could bind to the well-characterized acidic patch on the surface of nucleosomes. Instead of using mononucleosomes, we used the histone H2A-H2B heterodimer (41), which is a well-established surrogate for the acidic patch in NMR titration experiments with Sds3 CTD. Once again, no discernible perturbations could be detected in the NMR spectra, ruling out a potential role for the capped Tudor domain in nucleosome binding.

**Figure 3.**
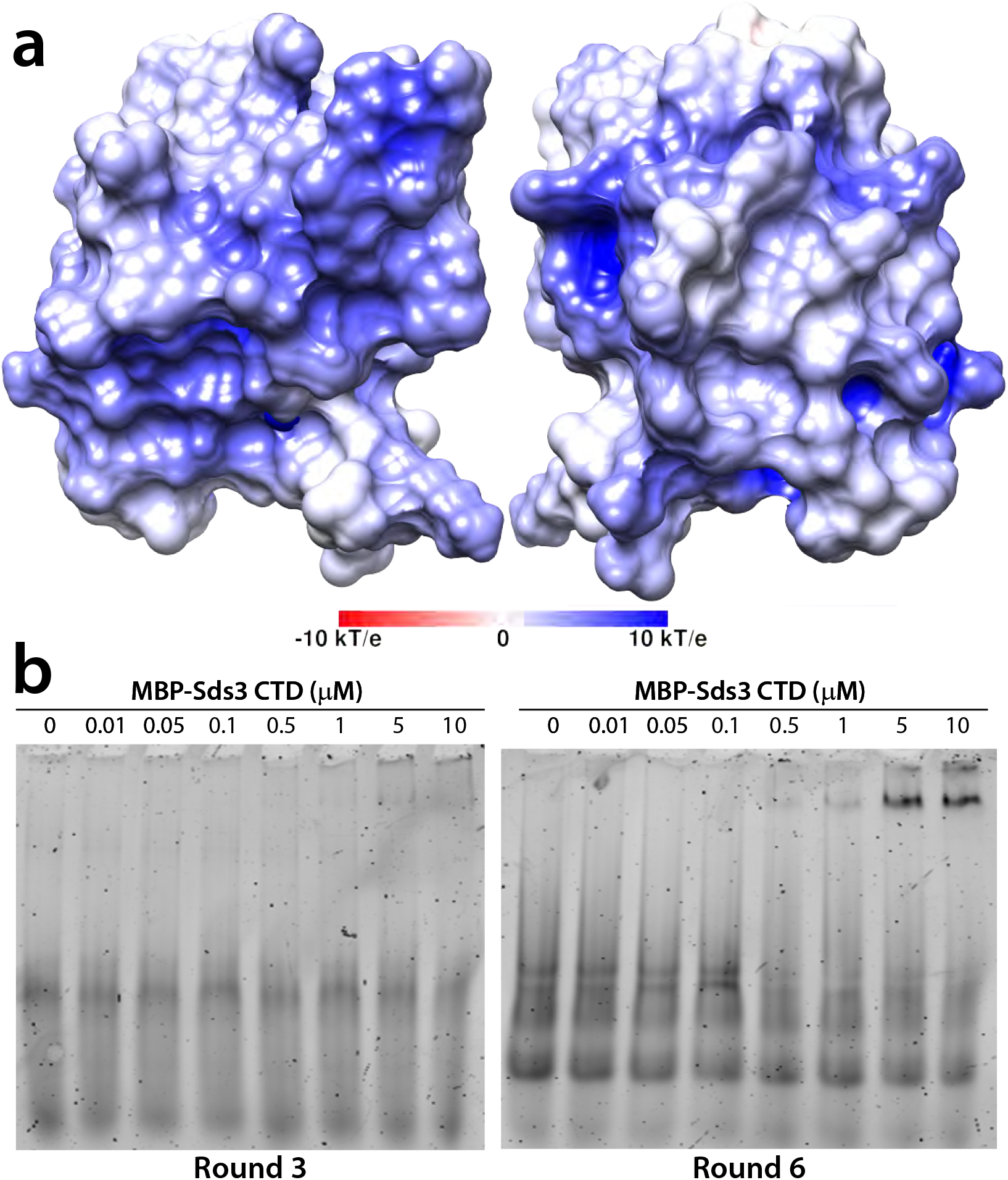
Analysis of the surface properties of Sds3 CTD suggests a nucleic acid-binding function. (**a**) An electrostatic potential map calculated using APBS and projected on to the molecular surface of Sds3 CTD. The poses for the two views are identical to those shown in figure 2. (**b**) Electrophoretic mobility shift assays conducted using MBP-tagged Sds3 CTD and the RNA library following round 3 (*left*) and round 6 (*right*) of the SELEX experiments. The library from round 6 was reverse transcribed and sent for NGS.

Since the knotted Tudor domain of Esa1 was previously shown to bind RNA (42), we asked whether the Sds3 CTD could have a similar function. To deduce potential RNA-binding motifs, SELEX experiments were performed starting with a 20mer randomized library (43). Samples of the RNA library following three rounds of selection and amplification were incubated with increasing amounts of MBP-tagged Sds3 CTD in electrophoretic mobility shift assays (EMSAs; **Figure 3b**). Although no clear mobility shifts were observed in these experiments, samples of the library following six rounds of selection, amplification, and incubation with MBP-Sds3 CTD yielded a clear band whose mobility was significantly reduced compared to that of the free RNA. These bands were observed at micromolar concentrations of MBP-Sds3 CTD, implying a modest affinity interaction. Following reverse transcription and amplification, the RNA library from this round was sent for next-generation sequencing (NGS). A total of 42 × 10^6^ reads were obtained, out of which ∼10 × 10^6^ were deemed to be of high quality for motif detection using the MEME suite. Because MEME can only handle a maximum of 500,000 sequences, the reads were randomly assigned to 10 datasets, each comprising 500,000 sequences. The five most statistically significant motifs in each dataset reported by MEME were compiled (**Supplementary Table S1**) and those that were found in more than three datasets are listed in **Table 2**. Somewhat unexpectedly and despite six rounds of enrichment, a single dominant motif did not emerge from these analyses. The most prevalent was a 7-residue motif (HGTGGTK; where H is A/C/T and K is G/T) found on average in 4.2% of the sequences. Remarkably, the other motifs deduced from these analyses were significantly enriched in Ts and Gs.

**Table 2.**
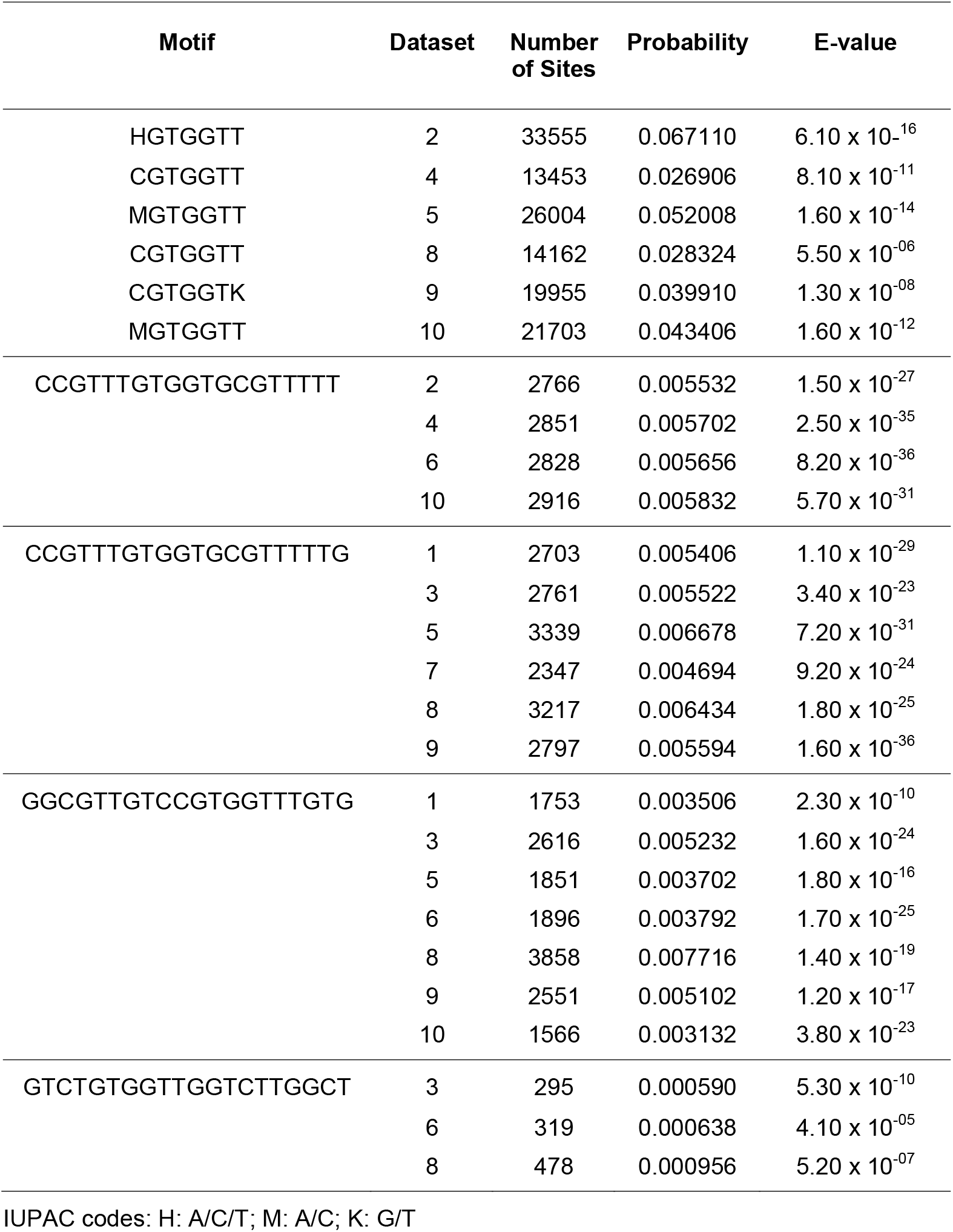
Statistically Significant Sequence Motifs Identified by MEME from NGS Data

### Sds3 CTD Binds G-quadruplexes

Since Ts and Gs are commonly found in G-quadruplexes, we asked whether a T_2_G_4_ DNA quadruplex might interact with Sds3 CTD (note that we chose to perform these experiments with DNA rather than RNA because both molecules can form similar quadruplex structures). We first recorded ^1^H NMR spectra to confirm G-quadruplex formation for this sequence, which as expected, is characterized by the presence of four narrow, imino proton resonances in the 10-11.5 ppm region emanating from each of the four tetrads ((44); **Supplementary Figure S3**). Four additional resonances of reduced intensity are observed in the imino proton region suggesting the formation of two types of quadruplexes that most likely differ in strand direction. The addition of one equivalent of T_2_G_4_ to ^15^N-Sds3 CTD induced significant perturbations in the NMR spectrum (**Figure 4a**) with a few resonances shifting to new positions and several others undergoing various degrees of broadening. Interestingly, the minor quadruplex species is relatively unperturbed in the presence of Sds3 CTD, implying that the interaction with the major species is specific (**Supplementary Figure S3**). Similarly, titrations with a random 10mer self-complementary DNA duplex (5’-GCGAATTCGC-3’) elicited only modest perturbations in the spectrum characterized by chemical shift deviations of 0.014 ± 0.012 ppm (**Supplementary Figure S4**). Additionally, virtually no perturbations were noted in the titration with a random RNA 8mer sequence (5’-AACUGUCG-3’). Collectively, these results indicate that Sds3 CTD preferentially associates with certain G-quadruplexes over double-stranded DNA or single-stranded RNA sequences.

**Figure 4.**
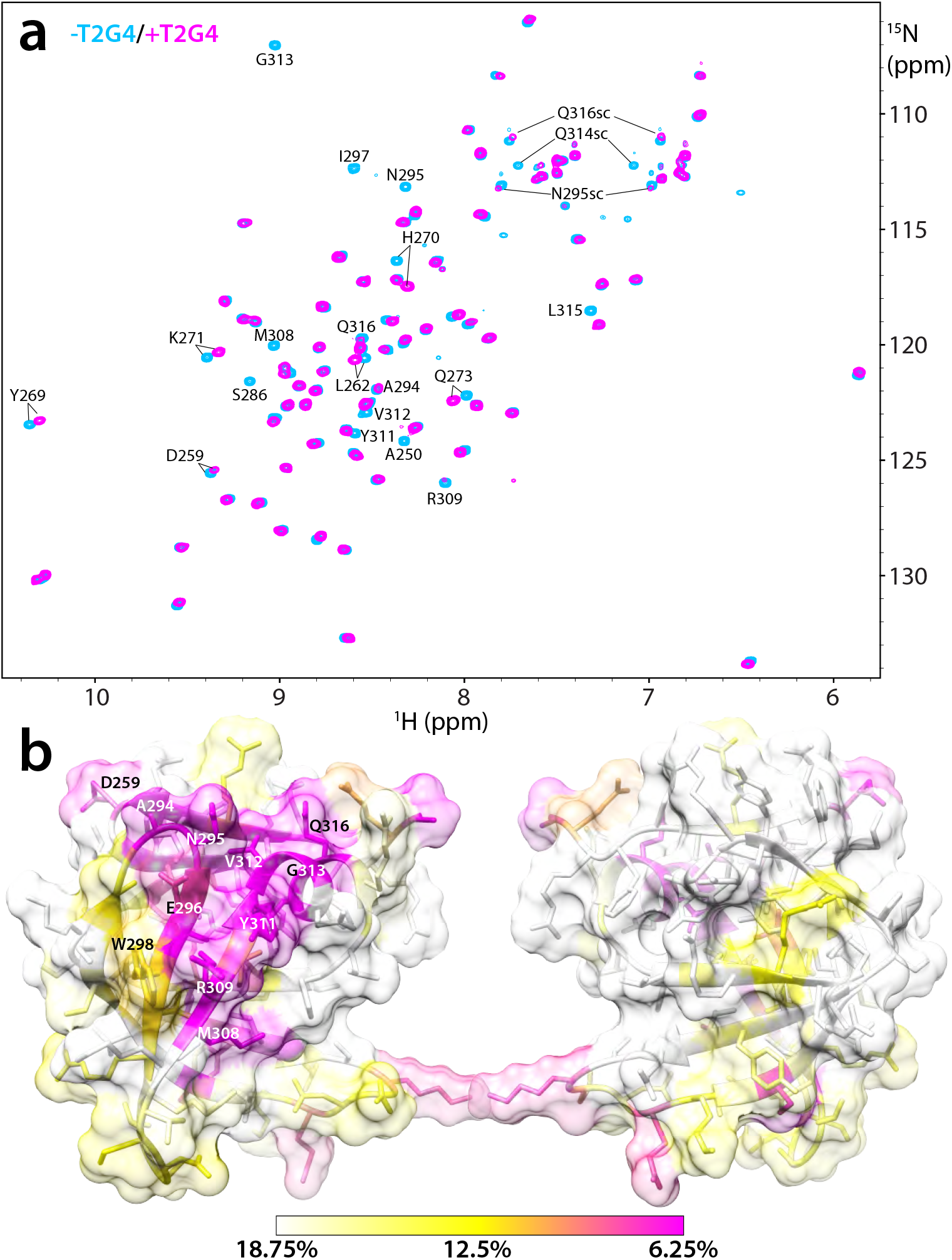
Sds3 CTD binds to a G-quadruplex. (**a**) ^1^H-^15^N HSQC spectra of Sds3 CTD recorded in the absence (blue) and presence (magenta) of 1 equivalent of T_2_G_4_ G-quadruplex DNA. Strongly perturbed resonances are annotated. To facilitate an objective comparison between the holo and apo spectra, the contour thresholds were adjusted using the peak intensities of the ‘unperturbed’ resonances. (**b**) Front and back views of the molecular surface of Sds3 CTD colored according to the ratio of the raw peak intensities in the holo and apo spectra (*i*.*e*., I_holo_/I_apo_). The surface is rendered semi-transparently to help identify the underlying residue. Note that the peak intensities of all resonances were diminished because of the larger size of the resulting complex and due to sample dilution caused by the addition of the quadruplex.

To identify the region of Sds3 CTD involved in binding to the G-quadruplex, we quantified the peak intensity ratios in the holo and apo HSQC spectra and mapped them on to the molecular surface of the CTD. The strongest perturbations were observed for a contiguous surface formed largely by residues in strands β6, β7, the loop preceding β6, and the sole helix connecting β7 and β8 (**Figure 4b**). This surface is distinct from the one located at the edge of the barrel that is commonly used by the BAH domain and the Royal family domains to engage with chromatin targets (**Figure 2a, 2b**, and **Supplementary Figure S2**). Interestingly, the three-stranded β-sheet formed by the capping motif of the CTD does not show significant spectral perturbations, implying that this novel feature is not essential for binding nucleic acids, at least those that were tested in this study.

## Discussion

HDAC containing chromatin-modifying complexes frequently contain many protein subunits with the non-enzymatic subunits widely thought to impart genome targeting specificity, especially since HDACs exhibit little sequence specificity themselves for acetylated targets. Two common mechanisms of genome targeting involve protein-protein interactions with sequence-specific DNA binding factors and/or engagement with specific post-translational modifications on histones. An especially intriguing feature of the Sin3L/Rpd3L complex is that the core subunits, including Sin3, Sds3, and SAP30, harbor domains of unknown structure and function that are narrowly distributed and found only in the respective orthologs and paralogs. The Sin3 subunit performs a scaffolding function for the assembly of the complex by engaging directly with most of the subunits while also providing multiple surfaces for direct engagement with DNA-bound factors. The SAP30 subunit is involved in turbocharging the catalytic activity of HDAC1 whereas Sds3 is thought to impart stability to the complex while also providing a dimerization function and interaction sites for DNA-bound factors and other subunits of the complex.

The discovery of a type of Tudor domain in Sds3 was unexpected and initially suggested an unrecognized function for the subunit in chromatin binding. However, as our subsequent studies have shown, Sds3 CTD shares more in common with another type of Tudor domain found in the Esa1 histone acetyltransferase previously shown to bind single-stranded RNA than with canonical Tudor domains (42). Although these non-canonical Tudor domains share a backbone RMSD of only 1.98 Å, with the highest level of structural similarity in the region spanning β4-β7 of Sds3 CTD, both domains bind to nucleic acids on the body of the barrel (**Figure 5**). The involvement of overlapping surfaces of the β-barrel in these distantly related domains is particularly striking. Even more interesting is the presence of conserved tryptophan and tyrosine residues at the protein-nucleic acid interface of both proteins (**Figure 5**), although the exact locations of these residues are not conserved between these domains. The latter likely reflects the different specificities of the domains for their target(s). Both the involvement of aromatic residues and their location on the surface of β-sheets are defining features shared with RNA-recognition motifs (45). Thus, both knotted and capped Tudor domains appear to have independently acquired nucleic acid-binding functions through a process of convergent evolution. Finally, since Esa1 is a histone acetyltransferase and a member of the NuA4 HAT complex (46), it is intriguing that these non-canonical Tudor domains are found in complexes with opposing enzymatic activities that produce contrasting transcriptional outcomes. Even more striking, the essential requirement for Esa1 in yeast can be bypassed through deletion of the gene encoding Sds3 (47).

**Figure 5.**
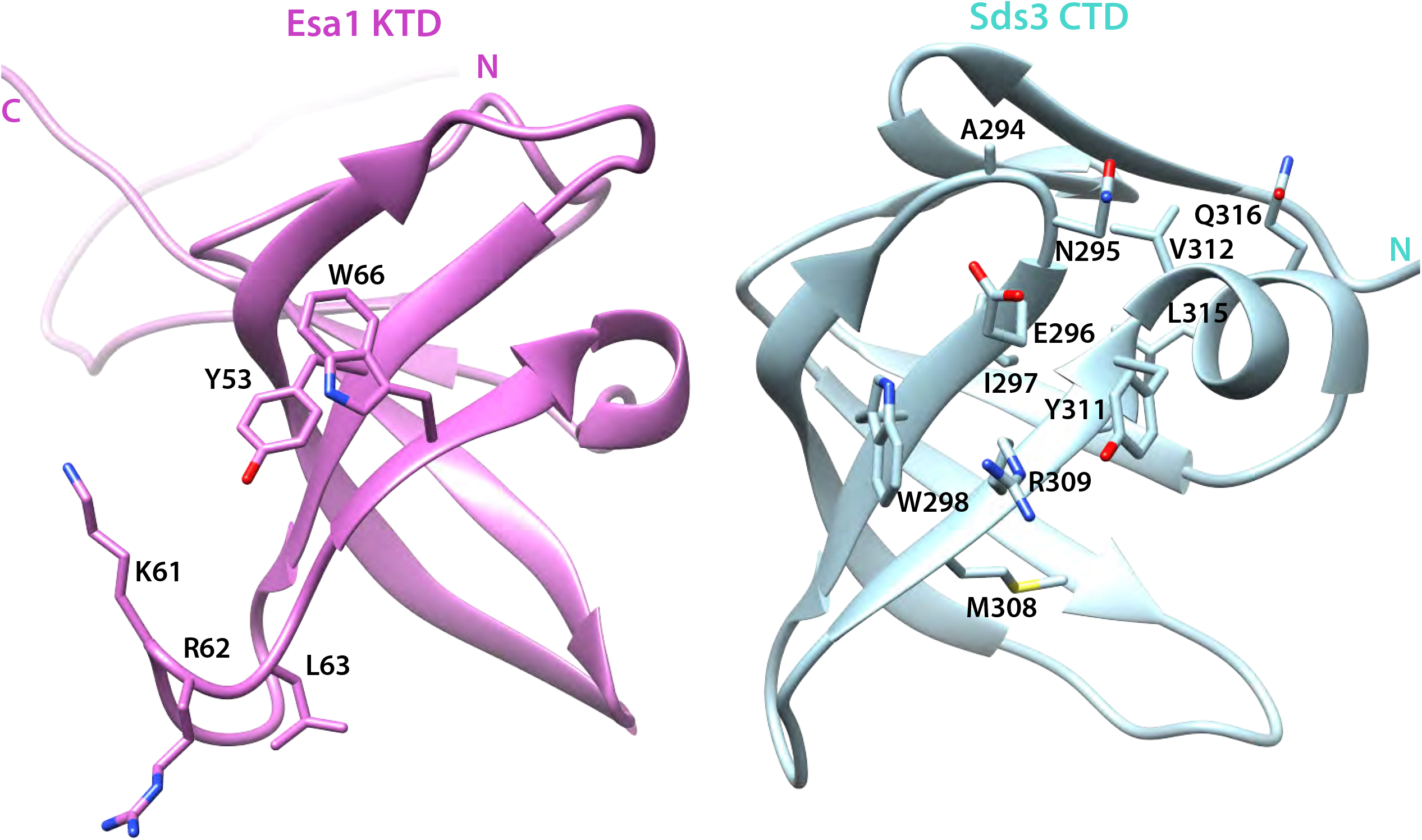
A side-by-side comparison following a best-fit backbone superposition of the Esa1 knotted Tudor domain (KTD; *left*; PDB ID: 2RO0)) with the Sds3 capped Tudor domain (*right*). The views highlight the residues that form the RNA and G-quadruplex binding surfaces inferred from NMR titration experiments.

There is precedent for RNA mediating the recruitment of at least one other chromatin-modifying complex, the LSD1-CoREST complex (48). The complex harbors both a histone demethylase (LSD1) as well as HDACs 1 and 2 and is recruited to the telomeric regions by a long non-coding RNA (lncRNA) that associates with chromatin in these regions to facilitate telomere silencing and heterochromatin formation (49). The lncRNA harbors many repeats of the 5’-UUAGGG-3’ sequence that forms intramolecular G-quadruplexes that in turn is critical for efficient interactions with LSD1 (50,51). Since G-quadruplexes have been found to localize to heterochromatin and gene knockout studies implicate Sds3 in the proper establishment of pericentric heterochromatin (16,52), it is tempting to speculate that the Sin3L/Rpd3L complex may be recruited to these regions through an analogous mechanism involving potentially G-quadruplex or other higher-order nucleic acid structures to promote gene silencing and heterochromatin formation at centromeres. A similar molecular mechanism may be operative in the context of R-loops where multiple subunits of the Sin3L/Rpd3L complex have been implicated in resolving these structures generated during transcription (25).

Although both Sds3 and BRMS1L share a CTD, the orthologous proteins define separate clades consistent with their distinct patterns of sequence conservation (**Figure 1a**), implying that the domains, while sharing a similar function, likely encode different specificities for their targets. Interestingly, the N-terminal three-stranded β-sheet that forms the capping motif that is unique to these Tudor domains does not seem to be involved in nucleic acid binding for the sequences that were tested. However, this structural motif also harbors conserved, solvent-exposed tyrosine and tryptophan residues (Y263 and W268) that could potentially be involved in binding other nucleic acid targets.

In conclusion, we have described a new type of Tudor domain that appears to have evolved to perform non-canonical functions such as binding nucleic acids. Although our results suggest that the Sds3 CTD has a preference for certain G-quadruplexes, more detailed studies focused on this domain are needed to definitively assess this preference over other higher-order nucleic acid structures as well as to identify the actual biologically relevant nucleic acid partner(s) targeted by the domain. If confirmed, it would only be the second instance of an HDAC-containing chromatin-modifying complex implicated in direct recruitment by higher-order nucleic acid structures, expanding the repertoire of macromolecules that could function in this manner. Our findings thus draw attention to potentially new and underappreciated roles for both Sds3 and the Sin3L/Rpd3L HDAC complex in transcription biology.

## Acknowledgements

This work was supported by grants from the American Heart Association to I.R. (17GRNT33680167) and R.D.M. (16PRE27260041) and from the National Institutes of Health for upgrading the 600 MHz NMR console (S10 OD012016). We thank members of the Radhakrishnan lab for critical comments. We are grateful to the Robert H. Lurie Comprehensive Cancer Center at Northwestern for supporting structural biology research. We thank Yawen Bai at the NCI for providing the plasmid encoding histone H2A-H2B heterodimer.

## Experimental Methods

### Construct Generation, Protein Expression and Purification

The coding sequence of mouse Sds3 CTD (residues 250-326) was sub-cloned into the pMCSG7, pMCSG9, and pMCSG10 bacterial expression vectors (53). His_6_-tagged-CTD encoded by the pMCSG7 vector was expressed at 16 °C in BL21(DE3) cells and subsequently purified via Ni^2+^-affinity chromatography. Cell pellets were resuspended in lysis buffer (50 mM Tris, pH 8.0, 200 mM NaCl, 1 mM TCEP, 1 mM phenylmethylsulfonyl fluoride (PMSF), 1 µM leupeptin, 1 mM pepstatin and 0.1% Triton X-100) and lysed by sonication. After centrifugation, the lysate was loaded onto a Ni^2+^-affinity resin (Sigma), washed with high salt (800 mM NaCl) and eluted with 300 mM imidazole. The His_6_-tag was removed by incubating the protein with TEV protease overnight at 4 °C, the samples concentrated and further purified via size exclusion chromatography using a Superdex 75 GL column (GE Healthcare) and a running buffer comprising 20 mM Tris (pH 8.0), 200 mM NaCl, and 1 mM DTT. Uniformly ^15^N- and/or ^13^C-labeled proteins were produced following the same procedure except they were grown in M9 minimal media supplemented with ^15^N-ammonium sulfate and/or ^13^C-glucose.

MBP- and GST-tagged proteins encoded by the pMCSG9 and pMCSG10 vectors were expressed and purified in a similar manner as the His_6_-tagged Sds3 CTD with the following changes. GST-Sds3 CTD was purified with glutathione sepharose (GE Healthcare) and eluted using 25 mM glutathione, while MBP-Sds3 was purified with amylose resin (New England Biolabs) and eluted with 25 mM maltose. All proteins were stored at 4 °C until they were used.

### NMR Spectroscopy and Structure Determination

All NMR spectra were acquired at 25 °C on a 600 MHz Agilent DD2 spectrometer. Sds3 CTD samples in the range of 350 *μ*M in 20 mM sodium phosphate buffer (pH 6.0) containing 50 mM NaCl, 1 mM DTT and 10% D_2_O were used to acquire NMR data. 3D HNCACB, CBCA(CO)NH, C(CO)NH-TOCSY, HNCO, and ^15^N-NOESY-HSQC NMR spectra were acquired for sequence-specific backbone resonance assignments (54). Data processing was performed using Felix (Felix NMR) and the peaks in these spectra were picked in NMRFAM-Sparky (55) and submitted to I-PINE for peak assignment (56). All the assignments were checked manually for accuracy. Side chain assignments were performed manually using 3D HCCH-COSY and HCCH-TOCSY spectra acquired in D_2_O; the sample for these experiments was generated by exchanging the buffer from H_2_O to D_2_O. Aromatic resonances were assigned based on a careful analysis of 2D ^1^H-^13^C aromatic HSQC, (HB)CB(CGCD)HD, (HB)CB(CGCDCE)HE (57) and ^1^H-^1^H NOESY spectra recorded in D_2_O.

Backbone ϕ and ψ dihedral angle restraints for structure calculations were derived from a combined analysis of the ^1^H^α, 13^C^α, 13^C^β, 13^C’, and backbone ^15^N chemical shifts using TALOS+ (58); only residues with reliability scores of 10 in secondary structural elements were restrained. ^1^H-^1^H NOE-based distance restraints were derived from three spectra, including 3D ^15^N-edited NOESY (τ_m_ = 80 ms) recorded in H_2_O, 3D ^13^C-edited aliphatic NOESY (τ_m_ = 60 ms), and 2D ^1^H-^1^H NOESY (τ_m_ = 75 ms) recorded in D_2_O.

Structures were determined using ARIA 1.2 in conjunction with CNS 1.1 starting from an initial structure with extended backbone conformation (59-61). All NOEs were calibrated automatically and were assigned iteratively by ARIA; the assignments were checked manually for errors after each run. Eighty conformers were calculated; 40 conformers with the lowest restraint energies were refined in a shell of water and the 20 conformers with the lowest restraint energies and violations and ideal covalent geometry were selected. The final conformers were analyzed using CNS (59), PROCHECK (62), and scripts written in-house.

### Histone Peptide Array

His_6_-tagged Sds3 CTD was incubated with a MODified™ histone peptide array (Active Motif) at a concentration of 15 μM. After a 2 h incubation period, the array was washed and probed with an anti-His primary antibody (Thermo Fisher, MA121315, 1:1000 dilution), after which anti-mouse HRP-conjugated secondary antibody (Thermo Fisher Scientific, #OB617005, 1:1000 dilution) was used. The array was imaged using West Pico chemiluminescent substrate (Thermo Scientific, #34080) and a Syngene Pxi chemiluminescent imager. The screen was performed in duplicate as a test of reproducibility.

### NMR Titrations

Sds3 CTD samples in the 150 to 230 *μ*M range in 20 mM sodium phosphate buffer (pH 6.0) containing 50 mM NaCl, 1 mM DTT, and 10% D_2_O were used for the NMR titration experiments. ^1^H-^15^N HSQC NMR spectra were acquired following the addition of excess dimethyllysine, trimethyllysine, acetyllysine, and dimethylarginine (Sigma; all compounds were used without further purification). NMR titrations were also conducted with an unmodified histone H3 peptide (residues 1-42), purified histone H2A-H2B heterodimer, DNA and RNA oligonucleotides. The oligonucleotides were purchased from Integrated DNA Technologies and Dharmacon and used without further purification. Data processing and analysis were performed using Felix (Felix NMR) and NMRFAM-Sparky (55).

### SELEX

SELEX (systematic evolution of ligands by exponential enrichment; (43) experiments were conducted with GST-tagged Sds3 CTD. An RNA library for selection experiments was obtained from TriLink Biotechnologies that consisted of random 20mer sequences flanked by adapters of known sequence for reverse transcription, PCR amplification, and sequencing. To preclear the RNA library of non-specific interactions with GST, the RNA library was initially incubated with purified GST immobilized on glutathione sepharose beads. The flow-through containing unbound RNA was then incubated with purified GST-Sds3 CTD immobilized on glutathione sepharose beads. The beads were washed extensively, and the protein was digested with proteinase K. Bound RNA was then purified using phenol/chloroform extraction. RNA was reverse transcribed and then PCR amplified. The PCR template was used to transcribe RNA and the process was repeated six times to enrich Sds3 CTD binding sequences.

After the final round of selection, PCR products were submitted to the Northwestern NUSeq core facility. Sequencing reads were generated using an Illumina SR75 sequencer. Sequences were trimmed to remove adapters using Cutadapt (63) and filtered by quality (Fast QC) in the Galaxy bioinformatics suite (64). Ten datasets of 500,000 randomly selected sequences from ∼10^6^ high quality reads were extracted using the Galaxy bioinformatics suite for motif analysis. Motifs were identified and analyzed using the MEME suite (65). The top five motifs from each dataset were compiled and those that appeared in more than three datasets were deemed significant for inclusion in Table 2.

### EMSAs

Samples were prepared by mixing increasing concentration of MBP-tagged Sds3 CTD with 1 *μ*M total RNA from rounds 3 and 6 (the final round) of SELEX. EMSAs were performed using 5% native PAGE gels with 0.5x TB buffer (50 mM Tris, 50 mM boric acid). Gels were equilibrated for 30 minutes before samples were loaded onto the gel. Samples were run at 4 °C for 90 min and then stained 30 min with SYBR™ Gold (ThermoFisher). Gels were imaged on a fluorescence Typhoon imager with the excitation and emission set to 480 nm and 520 nm, respectively.

## Abbreviations

BRMS1: breast cancer metastasis suppressor 1
BRMS1L: BRMS1-like
CTD: C-terminal domain/capped Tudor domain
HDAC: histone deacetylase
HAT: histone acetyltransferase
lncRNA: long non-coding RNA
MBP: maltose-binding protein
NGS: next-generation sequencing
RMS: root-mean-square
RMSD: RMS deviation
SELEX: systematic evolution of ligands by exponential enrichment
EMSA: electrophoretic mobility shift assay

## References

1. Gardner, K. E., Allis, C. D., and Strahl, B. D. (2011) OPERating ON Chromatin, a Colorful Language where Context Matters. J. Mol. Biol. 409, 36–46

2. Bowman, G. D., and Poirier, M. G. (2015) Post-translational modifications of histones that influence nucleosome dynamics. Chem. Rev. 115, 2274–2295

3. Evertts, A. G., Zee, B. M., Dimaggio, P. A., Gonzales-Cope, M., Coller, H. A., and Garcia, B. A. (2013) Quantitative dynamics of the link between cellular metabolism and histone acetylation. J. Biol. Chem. 288, 12142–12151

4. Moser, M. A., Hagelkruys, A., and Seiser, C. (2014) Transcription and beyond: the role of mammalian class I lysine deacetylases. Chromosoma 123, 67–78

5. Yang, X. J., and Seto, E. (2008) The Rpd3/Hda1 family of lysine deacetylases: from bacteria and yeast to mice and men. Nat. Rev. Mol. Cell Biol. 9, 206–218

6. Hayakawa, T., and Nakayama, J. (2011) Physiological roles of class I HDAC complex and histone demethylase. Journal of Biomedicine and Biotechnology 2011, 129383

7. Xue, Y., Wong, J., Moreno, G. T., Young, M. K., Cote, J., and Wang, W. (1998) NURD, a novel complex with both ATP-dependent chromatin-remodeling and histone deacetylase activities. Mol. Cell 2, 851–861

8. Smith, K. T., Sardiu, M. E., Martin-Brown, S. A., Seidel, C., Mushegian, A., Egidy, R., Florens, L., Washburn, M. P., and Workman, J. L. (2012) Human family with sequence similarity 60 member A (FAM60A) protein: a new subunit of the Sin3 deacetylase complex. Mol. Cell. Proteomics 11, 1815–1828

9. You, A., Tong, J. K., Grozinger, C. M., and Schreiber, S. L. (2001) CoREST is an integral component of the CoREST-human histone deacetylase complex. Proc. Natl. Acad. Sci. U. S. A. 98, 1454–1458

10. Bantscheff, M., Hopf, C., Savitski, M. M., Dittmann, A., Grandi, P., Michon, A. M., Schlegl, J., Abraham, Y., Becher, I., Bergamini, G., Boesche, M., Delling, M., Dumpelfeld, B., Eberhard, D., Huthmacher, C., Mathieson, T., Poeckel, D., Reader, V., Strunk, K., Sweetman, G., Kruse, U., Neubauer, G., Ramsden, N. G., and Drewes, G. (2011) Chemoproteomics profiling of HDAC inhibitors reveals selective targeting of HDAC complexes. Nat. Biotechnol. 29, 255–265

11. Guenther, M. G., Barak, O., and Lazar, M. A. (2001) The SMRT and N-CoR corepressors are activating cofactors for histone deacetylase 3. Mol. Cell. Biol. 21, 6091–6101

12. Carrozza, M. J., Florens, L., Swanson, S. K., Shia, W. J., Anderson, S., Yates, J., Washburn, M. P., and Workman, J. L. (2005) Stable incorporation of sequence specific repressors Ash1 and Ume6 into the Rpd3L complex. Biochim. Biophys. Acta 1731, 77-87; discussion 75-76

13. Silverstein, R. A., and Ekwall, K. (2005) Sin3: a flexible regulator of global gene expression and genome stability. Curr. Genet. 47, 1–17

14. Grzenda, A., Lomberk, G., Zhang, J. S., and Urrutia, R. (2009) Sin3: master scaffold and transcriptional corepressor. Biochim. Biophys. Acta 1789, 443–450

15. Adams, G. E., Chandru, A., and Cowley, S. M. (2018) Co-repressor, co-activator and general transcription factor: the many faces of the Sin3 histone deacetylase (HDAC) complex. Biochem. J. 475, 3921–3932

16. David, G., Turner, G. M., Yao, Y., Protopopov, A., and DePinho, R. A. (2003) mSin3-associated protein, mSds3, is essential for pericentric heterochromatin formation and chromosome segregation in mammalian cells. Genes Dev. 17, 2396–2405

17. Fleischer, T. C., Yun, U. J., and Ayer, D. E. (2003) Identification and characterization of three new components of the mSin3A corepressor complex. Mol. Cell. Biol. 23, 3456–3467

18. Banks, C. A. S., Thornton, J. L., Eubanks, C. G., Adams, M. K., Miah, S., Boanca, G., Liu, X., Katt, M. L., Parmely, T. J., Florens, L., and Washburn, M. P. (2018) A Structured Workflow for Mapping Human Sin3 Histone Deacetylase Complex Interactions Using Halo-MudPIT Affinity-Purification Mass Spectrometry. Mol. Cell. Proteomics 17, 1432–1447

19. Banks, C. A. S., Zhang, Y., Miah, S., Hao, Y., Adams, M. K., Wen, Z., Thornton, J. L., Florens, L., and Washburn, M. P. (2020) Integrative Modeling of a Sin3/HDAC Complex Sub-structure. Cell Rep 31, 107516

20. He, Y., Imhoff, R., Sahu, A., and Radhakrishnan, I. (2009) Solution structure of a novel zinc finger motif in the SAP30 polypeptide of the Sin3 corepressor complex and its potential role in nucleic acid recognition. Nucleic Acids Res. 37, 2142–2152

21. Marcum, R. D., and Radhakrishnan, I. (2019) Inositol phosphates and core subunits of the Sin3L/Rpd3L histone deacetylase (HDAC) complex up-regulate deacetylase activity. J. Biol. Chem. 294, 13928–13938

22. Clark, M. D., Marcum, R., Graveline, R., Chan, C. W., Xie, T., Chen, Z., Ding, Y., Zhang, Y., Mondragon, A., David, G., and Radhakrishnan, I. (2015) Structural insights into the assembly of the histone deacetylase-associated Sin3L/Rpd3L corepressor complex. Proc. Natl. Acad. Sci. U. S. A. 112, E3669–3678

23. Shi, X., Seldin, D. C., and Garry, D. J. (2012) Foxk1 recruits the Sds3 complex and represses gene expression in myogenic progenitors. Biochem. J. 446, 349–357

24. Meehan, W. J., Samant, R. S., Hopper, J. E., Carrozza, M. J., Shevde, L. A., Workman, J. L., Eckert, K. A., Verderame, M. F., and Welch, D. R. (2004) Breast cancer metastasis suppressor 1 (BRMS1) forms complexes with retinoblastoma-binding protein 1 (RBP1) and the mSin3 histone deacetylase complex and represses transcription. J. Biol. Chem. 279, 1562–1569

25. Salas-Armenteros, I., Perez-Calero, C., Bayona-Feliu, A., Tumini, E., Luna, R., and Aguilera, A. (2017) Human THO-Sin3A interaction reveals new mechanisms to prevent R-loops that cause genome instability. EMBO J. 36, 3532–3547

26. Nikolaev, A. Y., Papanikolaou, N. A., Li, M., Qin, J., and Gu, W. (2004) Identification of a novel BRMS1-homologue protein p40 as a component of the mSin3A/p33(ING1b)/HDAC1 deacetylase complex. Biochem. Biophys. Res. Commun. 323, 1216–1222

27. Gong, C., Qu, S., Lv, X. B., Liu, B., Tan, W., Nie, Y., Su, F., Liu, Q., Yao, H., and Song, E. (2014) BRMS1L suppresses breast cancer metastasis by inducing epigenetic silence of FZD10. Nature communications 5, 5406

28. Zhang, S., Lin, Q. D., and Di, W. (2006) Suppression of human ovarian carcinoma metastasis by the metastasis-suppressor gene, BRMS1. Int. J. Gynecol. Cancer 16, 522–531

29. Seraj, M. J., Samant, R. S., Verderame, M. F., and Welch, D. R. (2000) Functional evidence for a novel human breast carcinoma metastasis suppressor, BRMS1, encoded at chromosome 11q13. Cancer Res. 60, 2764–2769

30. Hurst, D. R. (2012) Metastasis suppression by BRMS1 associated with SIN3 chromatin remodeling complexes. Cancer Metastasis Rev. 31, 641–651

31. Smith, P. W., Liu, Y., Siefert, S. A., Moskaluk, C. A., Petroni, G. R., and Jones, D. R. (2009) Breast cancer metastasis suppressor 1 (BRMS1) suppresses metastasis and correlates with improved patient survival in non-small cell lung cancer. Cancer Lett. 276, 196–203

32. Silveira, A. C., Hurst, D. R., Vaidya, K. S., Ayer, D. E., and Welch, D. R. (2009) Over-expression of the BRMS1 family member SUDS3 does not suppress metastasis of human cancer cells. Cancer Lett. 276, 32–37

33. Hurst, D. R., and Welch, D. R. (2011) Unraveling the enigmatic complexities of BRMS1-mediated metastasis suppression. FEBS Lett. 585, 3185–3190

34. Zimmermann, R. C., and Welch, D. R. (2020) BRMS1: a multifunctional signaling molecule in metastasis. Cancer Metastasis Rev. 39, 755–768

35. Yang, J., Anishchenko, I., Park, H., Peng, Z., Ovchinnikov, S., and Baker, D. (2020) Improved protein structure prediction using predicted interresidue orientations. Proc. Natl. Acad. Sci. U. S. A. 117, 1496–1503

36. Holm, L. (2020) Using Dali for Protein Structure Comparison. Methods Mol. Biol. 2112, 29–42

37. Du, J., Zhong, X., Bernatavichute, Y. V., Stroud, H., Feng, S., Caro, E., Vashisht, A. A., Terragni, J., Chin, H. G., Tu, A., Hetzel, J., Wohlschlegel, J. A., Pradhan, S., Patel, D. J., and Jacobsen, S. E. (2012) Dual binding of chromomethylase domains to H3K9me2-containing nucleosomes directs DNA methylation in plants. Cell 151, 167–180

38. Xu, C., Cui, G., Botuyan, M. V., and Mer, G. (2015) Methyllysine Recognition by the Royal Family Modules: Chromo, Tudor, MBT, Chromo Barrel, and PWWP Domains.. in Histone Recognition (Zhou, M. M. ed.), Springer. pp 49–82

39. Jurrus, E., Engel, D., Star, K., Monson, K., Brandi, J., Felberg, L. E., Brookes, D. H., Wilson, L., Chen, J., Liles, K., Chun, M., Li, P., Gohara, D. W., Dolinsky, T., Konecny, R., Koes, D. R., Nielsen, J. E., Head-Gordon, T., Geng, W., Krasny, R., Wei, G. W., Holst, M. J., McCammon, J. A., and Baker, N. A. (2018) Improvements to the APBS biomolecular solvation software suite. Protein Sci. 27, 112–128

40. Vermeulen, M., Carrozza, M. J., Lasonder, E., Workman, J. L., Logie, C., and Stunnenberg, H. G. (2004) In vitro targeting reveals intrinsic histone tail specificity of the Sin3/histone deacetylase and N-CoR/SMRT corepressor complexes. Mol. Cell. Biol. 24, 2364–2372

41. Zhou, Z., Feng, H., Hansen, D. F., Kato, H., Luk, E., Freedberg, D. I., Kay, L. E., Wu, C., and Bai, Y. (2008) NMR structure of chaperone Chz1 complexed with histones H2A.Z-H2B. Nat. Struct. Mol. Biol. 15, 868–869

42. Shimojo, H., Sano, N., Moriwaki, Y., Okuda, M., Horikoshi, M., and Nishimura, Y. (2008) Novel structural and functional mode of a knot essential for RNA binding activity of the Esa1 presumed chromodomain. J. Mol. Biol. 378, 987–1001

43. Manley, J. L. (2013) SELEX to identify protein-binding sites on RNA. Cold Spring Harb Protoc 2013, 156–163

44. Adrian, M., Heddi, B., and Phan, A. T. (2012) NMR spectroscopy of G-quadruplexes. Methods 57, 11–24

45. Maris, C., Dominguez, C., and Allain, F. H. (2005) The RNA recognition motif, a plastic RNA-binding platform to regulate post-transcriptional gene expression. FEBS J. 272, 2118–2131

46. Avvakumov, N., and Cote, J. (2007) The MYST family of histone acetyltransferases and their intimate links to cancer. Oncogene 26, 5395–5407

47. Torres-Machorro, A. L., and Pillus, L. (2014) Bypassing the requirement for an essential MYST acetyltransferase. Genetics 197, 851–863

48. Khalil, A. M., Guttman, M., Huarte, M., Garber, M., Raj, A., Rivea Morales, D., Thomas, K., Presser, A., Bernstein, B. E., van Oudenaarden, A., Regev, A., Lander, E. S., and Rinn, J. L. (2009) Many human large intergenic noncoding RNAs associate with chromatin-modifying complexes and affect gene expression. Proc. Natl. Acad. Sci. U. S. A. 106, 11667–11672

49. Porro, A., Feuerhahn, S., and Lingner, J. (2014) TERRA-reinforced association of LSD1 with MRE11 promotes processing of uncapped telomeres. Cell Rep 6, 765–776

50. Patel, D. J., Phan, A. T., and Kuryavyi, V. (2007) Human telomere, oncogenic promoter and 5’-UTR G-quadruplexes: diverse higher order DNA and RNA targets for cancer therapeutics. Nucleic Acids Res. 35, 7429–7455

51. Hirschi, A., Martin, W. J., Luka, Z., Loukachevitch, L. V., and Reiter, N. J. (2016) G-quadruplex RNA binding and recognition by the lysine-specific histone demethylase-1 enzyme. RNA 22, 1250–1260

52. Hoffmann, R. F., Moshkin, Y. M., Mouton, S., Grzeschik, N. A., Kalicharan, R. D., Kuipers, J., Wolters, A. H., Nishida, K., Romashchenko, A. V., Postberg, J., Lipps, H., Berezikov, E., Sibon, O. C., Giepmans, B. N., and Lansdorp, P. M. (2016) Guanine quadruplex structures localize to heterochromatin. Nucleic Acids Res. 44, 152–163

53. Eschenfeldt, W. H., Lucy, S., Millard, C. S., Joachimiak, A., and Mark, I. D. (2009) A family of LIC vectors for high-throughput cloning and purification of proteins. Methods in molecular biology 498, 105–115

54. Ferentz, A. E., and Wagner, G. (2000) NMR spectroscopy: a multifaceted approach to macromolecular structure. Q Rev Biophys 33, 29–65

55. Lee, W., Tonelli, M., and Markley, J. L. (2015) NMRFAM-SPARKY: enhanced software for biomolecular NMR spectroscopy. Bioinformatics 31, 1325–1327

56. Lee, W., Bahrami, A., Dashti, H. T., Eghbalnia, H. R., Tonelli, M., Westler, W. M., and Markley, J. L. (2019) I-PINE web server: an integrative probabilistic NMR assignment system for proteins. Journal of biomolecular NMR 73, 213–222

57. Lohr, F., and Ruterjans, H. (1996) Novel Pulse Sequences for the Resonance Assignment of Aromatic Side Chains in ^13^C-Labeled Proteins. J. Magn. Reson. B 112, 259–268

58. Shen, Y., Delaglio, F., Cornilescu, G., and Bax, A. (2009) TALOS+: a hybrid method for predicting protein backbone torsion angles from NMR chemical shifts. J. Biomol. NMR 44, 213–223

59. Brünger, A. T., Adams, P. D., Clore, G. M., DeLano, W. L., Gros, P., Grosse-Kunstleve, R. W., Jiang, J. S., Kuszewski, J., Nilges, M., Pannu, N. S., Read, R. J., Rice, L. M., Simonson, T., and Warren, G. L. (1998) Crystallography & NMR system: A new software suite for macromolecular structure determination. Acta. Crystallogr. D Biol. Crystallogr. 54, 905–921.

60. Linge, J. P., Habeck, M., Rieping, W., and Nilges, M. (2003) ARIA: automated NOE assignment and NMR structure calculation. Bioinformatics 19, 315–316.

61. Linge, J. P., Habeck, M., Rieping, W., and Nilges, M. (2004) Correction of spin diffusion during iterative automated NOE assignment. J. Magn. Reson. 167, 334–342

62. Laskowski, R. A., Rullmannn, J. A., MacArthur, M. W., Kaptein, R., and Thornton, J. M. (1996) AQUA and PROCHECK-NMR: programs for checking the quality of protein structures solved by NMR. J. Biomol. NMR 8, 477–486.

63. Martin, M. (2011) Cutadapt Removes Adapter Sequences From High-Throughput Sequencing Reads. EMBnet 17

64. Afgan, E., Baker, D., Batut, B., van den Beek, M., Bouvier, D., Cech, M., Chilton, J., Clements, D., Coraor, N., Gruning, B. A., Guerler, A., Hillman-Jackson, J., Hiltemann, S., Jalili, V., Rasche, H., Soranzo, N., Goecks, J., Taylor, J., Nekrutenko, A., and Blankenberg, D. (2018) The Galaxy platform for accessible, reproducible and collaborative biomedical analyses: 2018 update. Nucleic acids research 46, W537–W544

65. Bailey, T. L., Boden, M., Buske, F. A., Frith, M., Grant, C. E., Clementi, L., Ren, J., Li, W. W., and Noble, W. S. (2009) MEME SUITE: tools for motif discovery and searching. Nucleic acids research 37, W202–208

