## Supplementary Materials for "A Capped Tudor Domain within a Core Subunit of the Sin3L/Rpd3L Histone Deacetylase Complex Binds Nucleic Acids"

**Supplementary Table S1**

| Dataset | Motif | Sites | E-value | Probability |
|---|---|---|---|---|
| 1 | CCGTTTGTGGTGCGTTTTTG | 2703 | 1.10E-29 | 0.005406 |
| 1 | GGCGTTGTCCGTGGTTTGTG | 1753 | 2.30E-10 | 0.003506 |
| 1 | CAGTGTCKTGKTTGTG | 3793 | 1.40E-03 | 0.007586 |
| 1 | CTGCTTCGTTTTGCTTGTGG | 165 | 7.30E-04 | 0.00033 |
| 1 | CCGTDVGWTRGTTDGGCATG | 2407 | 7.90E-02 | 0.004814 |
| 2 | CCGTTTGTGGTGCGTTTTT | 2766 | 1.50E-27 | 0.005532 |
| 2 | HGTGGTT | 33555 | 6.10E-16 | 0.06711 |
| 2 | CGTCCGAATATGTGGATGCT | 57 | 1.40E-04 | 0.000114 |
| 2 | TGCGGTGTCCGTGGGCRTGA | 926 | 3.30E-03 | 0.001852 |
| 2 | CCDWARRDGTGKYGTG | 3197 | 4.70E-03 | 0.006394 |
| 3 | GGCGTTGTCCGTGGTTTGTG | 2616 | 1.60E-24 | 0.005232 |
| 3 | CCGTTTGTGGTGCGTTTTTG | 2761 | 3.40E-23 | 0.005522 |
| 3 | GTCTGTGGTTGGTCTTGGCT | 295 | 5.30E-10 | 0.00059 |
| 3 | TGCGTAYGTYGDTAGTGG | 3330 | 1.20E-08 | 0.00666 |
| 3 | CGTGGTTKT | 10498 | 9.70E-05 | 0.020996 |
| 4 | CCGTTTGTGGTGCGTTTTT | 2851 | 2.50E-35 | 0.005702 |
| 4 | CGTGGTT | 13453 | 8.10E-11 | 0.026906 |
| 4 | GGTTTTGGTCGCTACTACGA | 621 | 3.90E-08 | 0.001242 |
| 4 | STATGTAGAAAAGTTG | 2037 | 4.90E-05 | 0.004074 |
| 4 | CCGATTTGTTGGCA | 991 | 1.50E-01 | 0.001982 |
| 5 | CCGTTTGTGGTGCGTTTTTG | 3339 | 7.20E-31 | 0.006678 |
| 5 | GGCGTTGTCCGTGGTTTGTG | 1851 | 1.80E-16 | 0.003702 |
| 5 | MGTGGTT | 26004 | 1.60E-14 | 0.052008 |
| 5 | CGTCCATCGTGGCCATGAGA | 34 | 7.00E-06 | 0.000068 |
| 5 | AGTGTCTGTATTGTGTGGCA | 508 | 1.10E-01 | 0.001016 |
| 6 | CCGTTTGTGGTGCGTTTTT | 2828 | 8.20E-36 | 0.005656 |
| 6 | GGCGTTGTCCGTGGTTTGTG | 1896 | 1.70E-25 | 0.003792 |
| 6 | GTCTGTGGTTGGTCTTGGCT | 319 | 4.10E-05 | 0.000638 |
| 6 | CRTCCARMKMKGGTTSDTGC | 759 | 1.20E-04 | 0.001518 |
| 6 | WAAWGTGGC | 7650 | 4.10E-03 | 0.0153 |
| 7 | CCGTTTGTGGTGCGTTTTTG | 2347 | 9.20E-24 | 0.004694 |
| 7 | GGCGWTGTCCGTGGTKYRTG | 2649 | 1.80E-10 | 0.005298 |

| | | | | |
|---|---|---|---|---|
| 7 | CRCRGTGKYKYGTTTKWGGT | 2094 | 8.30E-07 | 0.004188 |
| 7 | CWRAKTGTGGTTKYGM | 3350 | 1.60E-06 | 0.0067 |
| 7 | CMWDMGTGG | 3626 | 1.30E-03 | 0.007252 |
| 8 | CCGTTTGTGGTGCGTTTTTG | 3217 | 1.80E-25 | 0.006434 |
| 8 | GGCGTTGTCCGTGGTTTGTG | 3858 | 1.40E-19 | 0.007716 |
| 8 | GTCTGTGGTTGGTCTTGGCT | 478 | 5.20E-07 | 0.000956 |
| 8 | CGTGGTT | 14162 | 5.50E-06 | 0.028324 |
| 8 | TGTGGCTTTGCGTGGTATG | 2 | 2.50E-03 | 0.000004 |
| 9 | CCGTTTGTGGTGCGTTTTTG | 2797 | 1.60E-36 | 0.005594 |
| 9 | GGCGTTGTCCGTGGTTTGTG | 2551 | 1.20E-17 | 0.005102 |
| 9 | GVVGYGTCCGTSMGTG | 5079 | 1.90E-15 | 0.010158 |
| 9 | TGMTRYGTMSAMAAGTTGKC | 585 | 2.30E-11 | 0.00117 |
| 9 | CGTGGTK | 19955 | 1.30E-08 | 0.03991 |
| 10 | CCGTTTGTGGTGCGTTTTT | 2916 | 5.70E-31 | 0.005832 |
| 10 | GGTGATGTCCGTGGAGCATG | 705 | 1.30E-26 | 0.00141 |
| 10 | GGCGTTGTCCGTGGTTTGTG | 1566 | 3.80E-23 | 0.003132 |
| 10 | MGTGGTT | 21703 | 1.60E-12 | 0.043406 |
| 10 | TGCGTGTCTTCAGGTAAGGC | 195 | 4.20E-09 | 0.00039 |

IUPAC codes: M: A/C; R: A/G; W: A/T; S: C/G; Y: C/T; K: G/T; V: A/C/G; H: A/C/T; D: A/G/T; B: C/G/T; N: A/G/C/T

### Supplementary Figures

**Supplementary Figure S1**. Sds3 CTD is a folded domain. $^1$H-$^{15}$N HSQC spectrum of the CTD recorded at 25 °C showing narrow, well-dispersed amide proton and nitrogen resonances. Sequence-specific and side chain assignments are annotated.

**Supplementary Figure S2**. Sds3 CTD shares the β-barrel fold with members of the Royal family domains including the Tudor, PWWP, MBT, and Chromobarrel domains. The overall fold of a representative member of each domain type is shown along with the PDB accession code. For ease of comparison, each domain was first superimposed onto the Sds3 CTD. In each case, methyllysine-bearing histone peptides engage with the same edge of the open barrel. Although details of the binding modes vary, a common theme shared by all members of the family is that a 'cage' formed by aromatic residues at one edge of these β-barrel domains serves as the receptor for methyllysine residues (shown in space-filling representation). The backbone and side chains of the CTD are shown in a comparable pose to the other domains to emphasize the general lack of aromatic residues along the edge of the barrel that engages with methyllysine residues.

**Supplementary Figure S3**. The $T_2G_4$ DNA sequence forms quadruplexes and interacts with Sds3 CTD. Expanded plots of the imino proton region of the 1D $^1$H NMR spectra of apo $T_2G_4$ (*top*) and in the presence of one and two equivalents of $^{15}$N-Sds3 CTD (*middle* and *bottom panels*, respectively). Perturbations of the imino proton resonances of the dominant quadruplex species are readily observed. A second quadruplex species is also observed (denoted by filled circles) that unlike the major species does not appear to be perturbed by protein binding. Note that the spectra were recorded without $^{15}$N decoupling, giving rise to resonances (denoted by asterisks) belonging to the protein that are split by the $^1J_{NH}$ coupling.

**Supplementary Figure S4**. Double-stranded DNA binds with low affinity and produces only modest perturbations in the NMR spectrum of Sds3 CTD when compared to those induced by the $T_2G_4$ G-quadruplex DNA in **Figure 4a**. (**a**) $^1$H-$^{15}$N HSQC spectra of Sds3 CTD recorded in the absence and presence of 1 equivalent of a self-complementary, double-stranded duplex 5'-GCGAATTCGC-3'. Resonances of those residues that are significantly perturbed are annotated. (**b**) Chemical shift perturbations (CSP = $\sqrt{}0.5*(\Delta\delta H^2 + (\Delta\delta N/5)^2)$ graphed as a function of residue number. The horizontal lines across the graph denote values corresponding to the average CSP (<CSP>) and 1 and 2 standard deviations (σ) above the average.

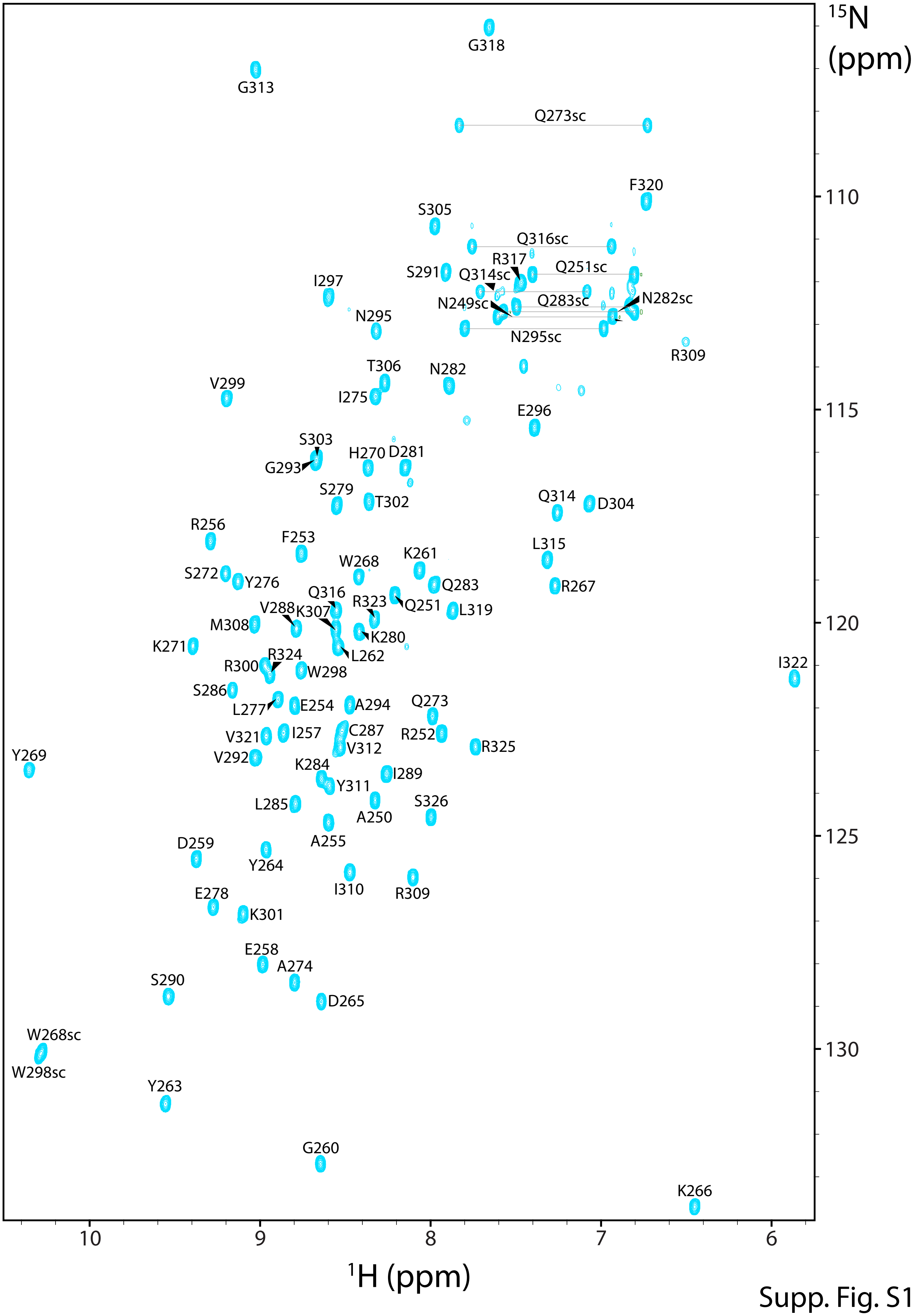
15N
(ppm)
110
115
120
125
130
10
9
8
7
6
1H (ppm)
G318
G313
Q273sc
F320
S305
Q316sc
R317
Q251sc
S291
Q314sc
I297
N249sc
Q283sc
N282sc
N295
N295sc
R309
T306
N282
V299
I275
E296
S303
H270
D281
G293
S279
T302
Q314
D304
R256
F253
L315
K261
W268
S272
Y276
Q283
R267
Q316
R323
Q251
V288
K307
L319
M308
K280
K271
R324
L262
R300
W298
I322
S286
Q273
E254
A294
L277
C287
R252
V321
I257
V312
R325
V292
Y269
K284
I289
Y311
L285
S326
A250
A255
D259
Y264
I310
R309
E278
K301
E258
A274
S290
D265
W268sc
W298sc
Y263
G260
K266

Supp. Fig. S1

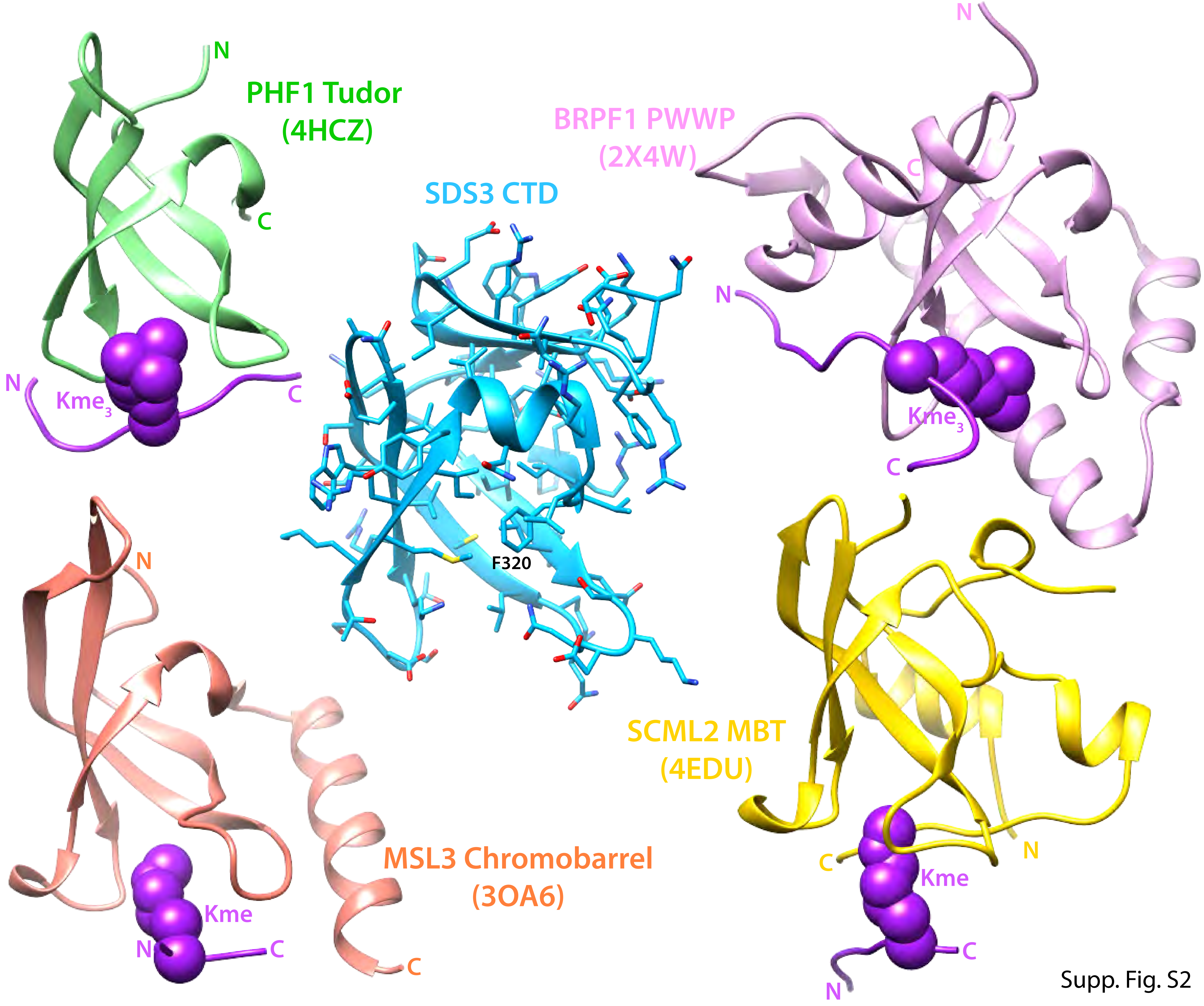

PHF1 Tudor
(4HCZ)
N
C
N
Kme$_3$
C
SDS3 CTD
F320
BRPF1 PWWP
(2X4W)
N
C
N
Kme$_3$
C
N
MSL3 Chromobarrel
(3OA6)
Kme
N
C
C
SCML2 MBT
(4EDU)
N
C
Kme
C
N

Supp. Fig. S2

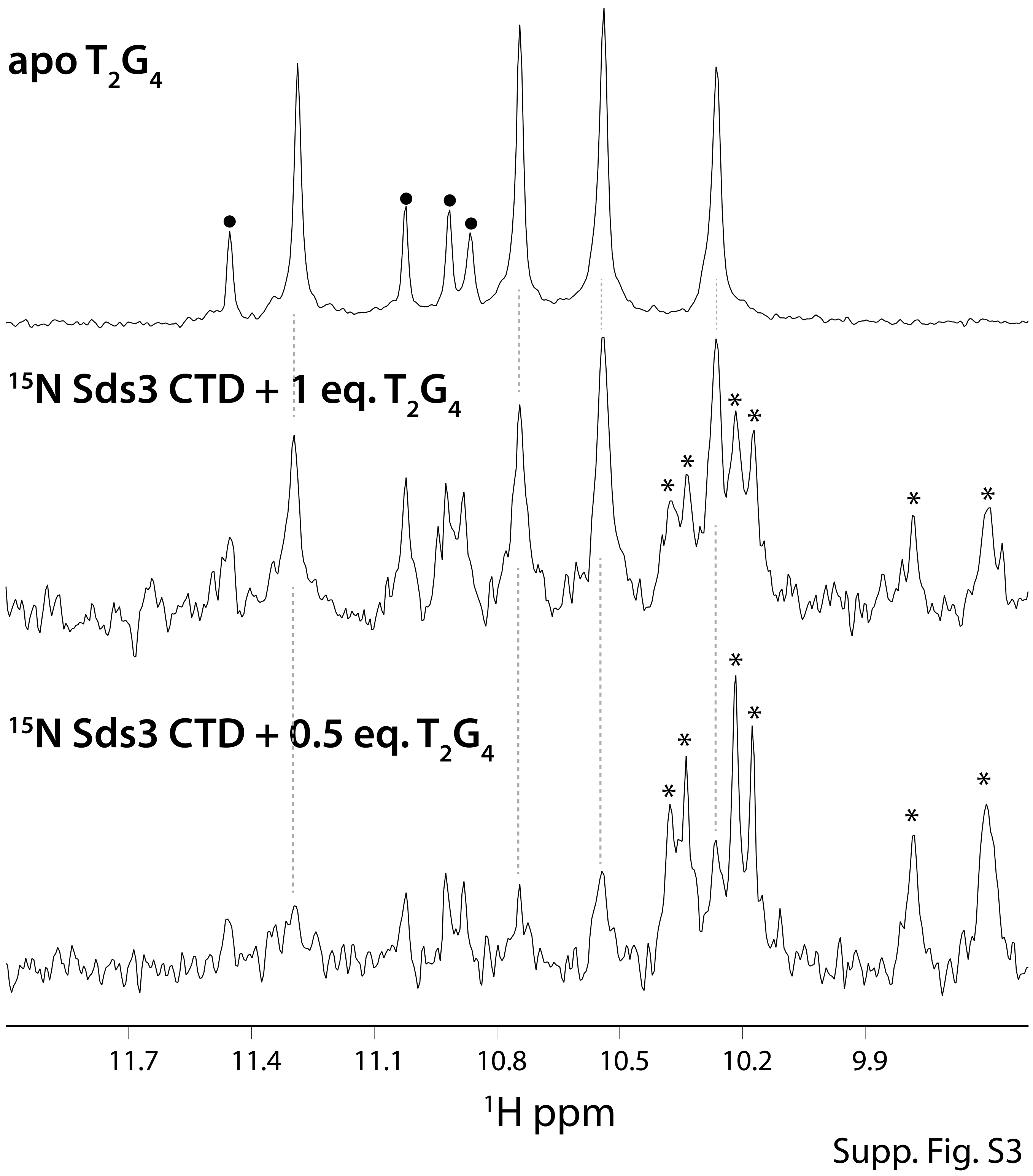
apo $T_2G_4$
$^{15}$N Sds3 CTD + 1 eq. $T_2G_4$
$^{15}$N Sds3 CTD + 0.5 eq. $T_2G_4$
11.7
11.4
11.1
10.8
10.5
10.2
9.9
$^1$H ppm

Supp. Fig. S3

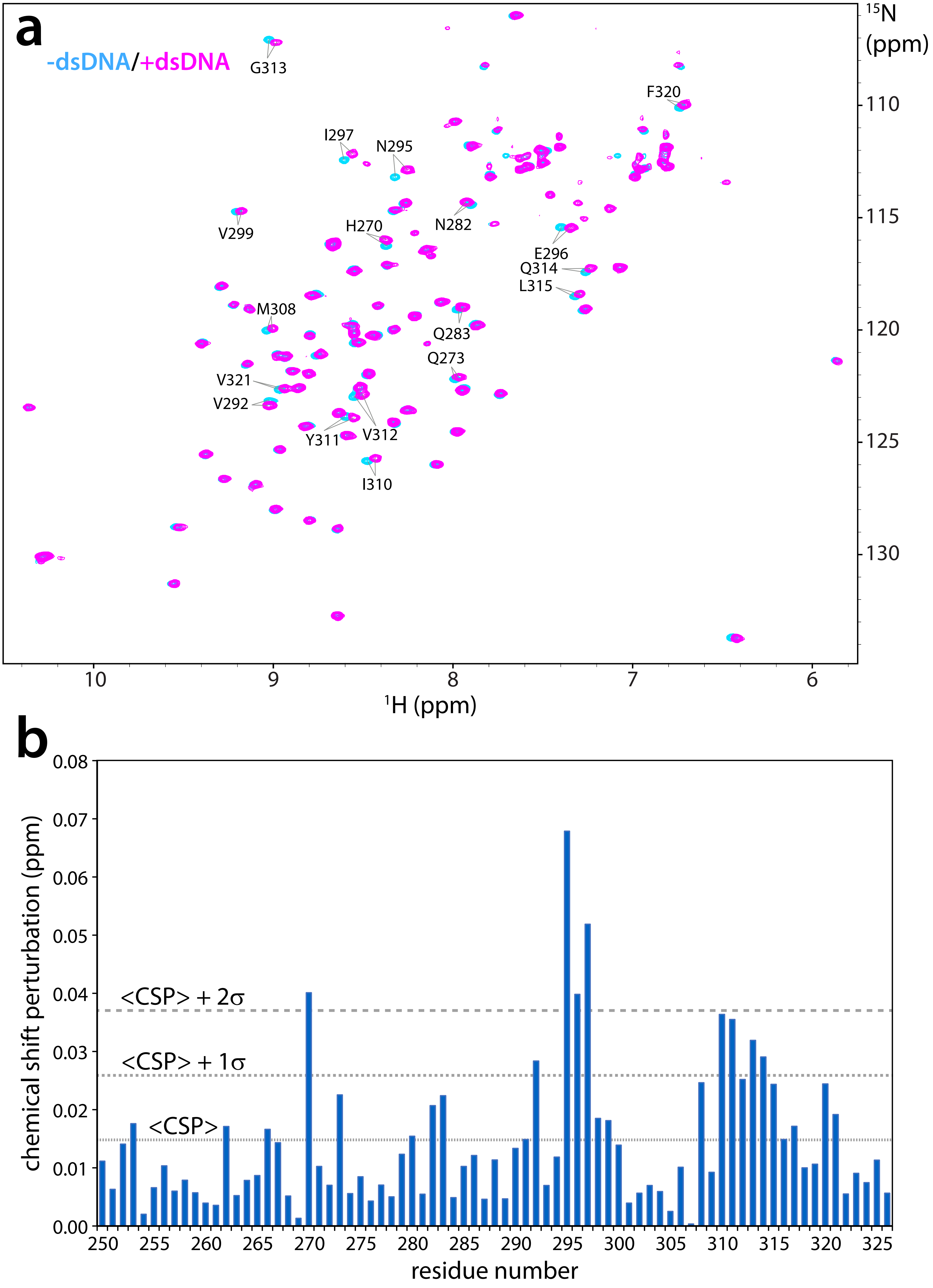
a
-dsDNA/+dsDNA
G313
F320
I297
N295
V299
H270
N282
E296
Q314
L315
M308
Q283
Q273
V321
V292
Y311
V312
I310
15N (ppm)
110
115
120
125
130
10
9
8
7
6
1H (ppm)
b
chemical shift perturbation (ppm)
0.08
0.07
0.06
0.05
0.04
0.03
0.02
0.01
0.00
<CSP> + 2σ
<CSP> + 1σ
<CSP>
250
255
260
265
270
275
280
285
290
295
300
305
310
315
320
325
residue number

Supp. Fig. S4
